# Effect of TSS on anti-CVB3 through TLR3 pathway

**DOI:** 10.1101/2022.03.28.486160

**Authors:** Xiao-han Zheng, Ming-ming Yuan, Yuan Wang, Jing Zhou, Lei Zhang

## Abstract

Toll-like receptors (TLRs) can recognize molecular patterns associated with microorganisms and induce immune responses. TLR3 is responsible for recognizing viruses and making hosts resistant to Coxsackie B3 (CVB3) infection. In this study, we found that Tectorigenin Sodium Sulfonate (TSS) has a TLR3 inhibitory effect and exerts strong anti-CVB3 activity through the TLR3 pathway both in vivo and in vitro. Balb/c mice were infected by ip injection of CVB3 virus, and subcutaneously injected with TSS at 264mg·Kg^-1^·d^-1^ for 7 d. The life extension rate of CVB3-infected mice was up to 46.67%, and the mean time to death was prolonged from 6.00 ± 0.47 d to 8.80 ± 2.78 d (*P < 0.01*). TSS was also able to reduce the level of weight loss in mice, significantly reduce the symptoms of myocarditis caused by CVB3, reduce the virus titer in the heart, improve the spleen and thymus index, and enhance immune function. We further explored the underlying mechanism of TSS in CVB3-infected HT-29 cells and mice: TSS reduced TLR3 expression and downregulated its downstream TRIF, TRAF6, IRF3, NF-κ B. MAPK, AP1 expression, reduced expression of IFN-β、IL-6、TNF-α. TSS has strong anti-CVB3 activity in vitro and in vivo and has the effect of inhibiting the activation of TLR3 pathway.

**IMPORTANCE:** TSS is expected to be an antiviral drug for the treatment of CVB3 infection. Its regulatory effect on the immune system provides a new approach for antiviral research.

## INTRODUCTION

Viral myocarditis (VMC) produces nonspecific interstitial myocardial inflammation and necrosis, leading to malignant arrhythmias, heart failure, and sudden cardiac death in patients^(1)^. A variety of viral infections can cause VMC, among which Coxsackievirus B3 (CVB3) is the most common and widely studied pathogen clinically^(2)^. Although VMC has been extensively studied, challenges remain in the diagnostic and treatment. Therefore, it is necessary to understand the pathogenesis of VMC and search for novel and effective drugs for VMC.

The viral replication mechanism of myocarditis induced by viral infection of cardiomyocytes is driven by viral infection-induced cell destruction and infection-triggered immune system activation, so the inflammatory process is critical for pathogenesis^(3)^. In addition to direct virus-induced cardiac cytotoxicity, the immune system and host genetics play a central role in myocardial inflammation^(4)^. An immune response is necessary to control the virus, but inflammatory processes lead to cardiac damage and fibrotic remodeling^(5)^. Toll-like receptors (TLRs) are a class of innate immune recognition receptors that recognize molecular patterns associated with microorganisms and induce immune responses^(6)^. DsRNA is a significant activator of the innate immune system, as part of the viral life cycle at the same time, is the heritable material of some types of viruses or the product of most viruses^(7, 8)^ and a typical pathogen-associated molecular pattern (PAMP) molecules, which can be recognized by TLR3^(9, 10)^. The role of TLR3 in viral protective immunity has been studied in mice and humans with genetically impaired TLR3 responses^(11)^, it has been found that play an important role in cardiovascular disease and viral myocarditis. TLR3 transmits risk signals to Nuclear factors-κB (NF-κB) through activation of TRIF (TIR domain-containing linker-induced interferon-κ), TRAF6 (tumor necrosis factor receptor-associated factor 6), and IRAK1 (interleukin 1 receptor-associated kinase 1),activated NF-κB leads to the translocation of its p50 and p65 subunits from the cytoplasm to the nucleus and drives the expression of proinflammatory cytokines^(12)^. TLR3 signaling initiates the innate immune response to the virus leading to the expression of IFN, which activates the Ifnb1 gene through p38 MAPK as well as AP-1, NF-κB and IRF-3^(13–15)^. In addition, TLR3-mediated p38MAPK activation can stably activate Ifnb1 mRNA expression^(16)^. IFN-I signaling is required for the innate immune response to CVB3 infection^(17)^, and mice lacking IFN-β are more susceptible to CVB3 infection than wild-type mice^(18)^, with reduced administration of IFN-α or IFN-β CVB3-induced myocarditis in mice and humans^(19–21)^.

Rhizoma Iridis Tectori is the dry rhizome of *Iris tectorum Maxim*., which has the functions of heat-clearing and detoxifying, eliminating phlegm, and soothing the throat^(22)^. Tectorigenin Sodium Sulfonate (TSS) is a new type of antiviral drug obtained by structural modification of flavonoid active ingredients in Rhizoma Iridis Tectori, which significantly increases the water solubility of irisin. Previous studies in our laboratory have shown that TSS has a certain degree of inhibitory effect on parainfluenza virus, SARS and enterovirus CVB3, but the mechanism remains unclear. This study further investigated the anti-CVB3 virus activity of TSS in vivo and in vitro, and carried out mechanism studies to explore its mode of action. Our findings suggest that TSS is a potential antiviral drug that can inhibit viral replication and regulate TLR3 pathway to regulate inflammation to exert antiviral effects.

## MATERIALS AND METHODS

### Viruses and cells

Coxsackievirus B3 strain (Nancy) was purchased from the China Center for Type Culture Collection. Virus amplification and preservation are carried out by the laboratory. The virus operations designed in this experiment are handled in the safety protection second-level laboratory.

Mouse lung cancer cells (LLC) and human colon cancer cells (HT-29) were cultured in Dulbecco’s Modified Eagle’s medium (DMEM) and DME/F-12 medium, respectively. The DMEM medium was supplemented with 10% fetal bovine serum (FBS), add 5% FBS to the DME/F-12 medium, and incubate at 37°C in a 5% CO_2_ incubator.

### Samples

Tectorigenin Sodium Sulfonate (TSS) was provided by Sichuan Academy of Traditional Chinese Medicine, is a light yellow powder with a purity of 99.8 %, publication (Bulletin) No: CN1285586C, protected from light tightly preserved for further use, the structure is shown in Fig. 1a; Guanidine hydrochloride (Ghc) standard was purchased from Beijing solarbio science & technology co.,ltd, with product lot 625G021 and CAS number 50-01-1; Ribavirin injection was purchased from Shiyao Yinhu Pharmaceutical Co., Ltd., specification 100mg/mL, lot: 1. 012102253.

**Fig. 1.**
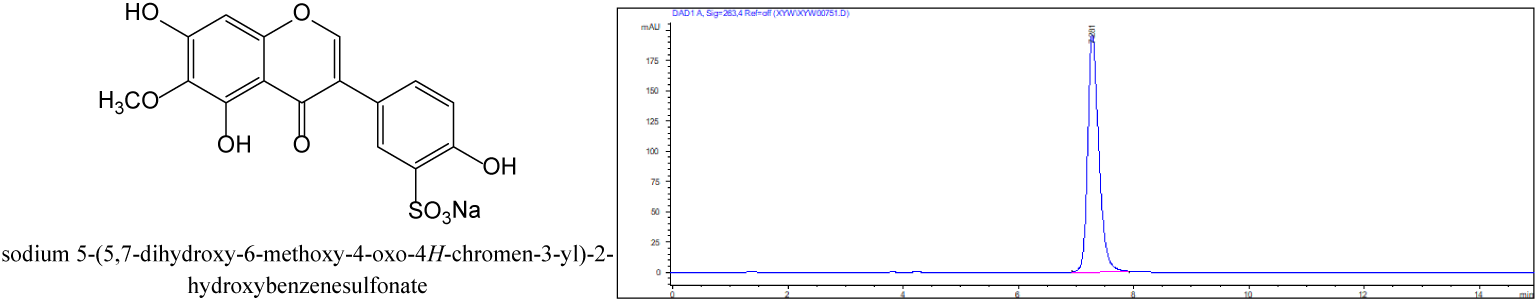
The chemical structure of TSS and the HPLC patterns of TSS

**Fig. 2.**
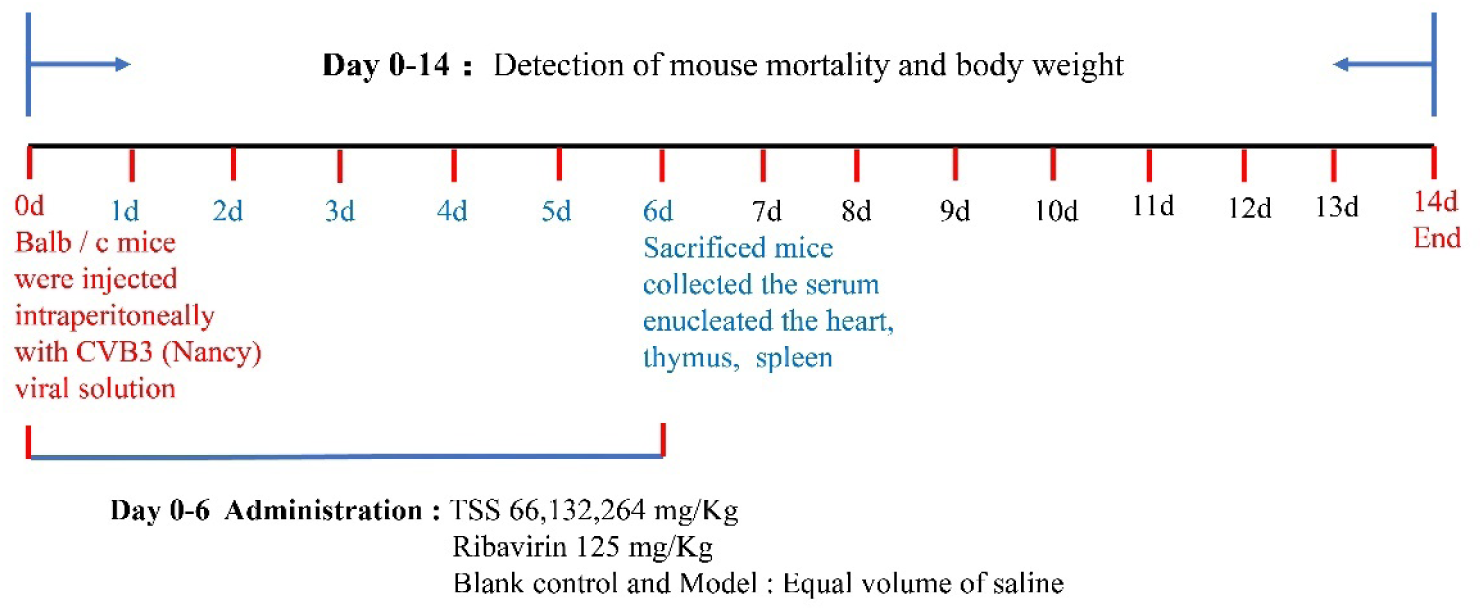
Basic experimental protocol for TSS treatment of CVB3 infection: mice were infected with CVB3 virus on day 0 (except the blank control group), and 3 h later, all mice were intraperitoneally injected with TSS (264, 132 and 66 mg·Kg^-1^·d^-1^), Ribavirin (125 mg·Kg^-1^·d^-1^), the blank control group and model group did not receive any treatment, were continuously administered for 6d(each group n=10).

### Mice

Male Balb/c mice, 3-4 weeks old, were purchased from Beijing Vital River Laboratory Animal Technology Co., Ltd, were housed in independent ventilated cages and received pathogen free food and water. All animals received care according to *The Guide for the Care and Use of Laboratory Animals*.

### Therapeutic efficacy study in mice

The CVB3 Nancy strain was diluted with normal saline to 350 TCID_50_ (to cause the mice to die) and 25 TCID_50_ (Cytokine expression was made significantly different between mice), and the mice were inoculated with 100 μL of virus fluid i.p. After 3 h, the administration group was intraperitoneally injected with TSS or ribavirin of different concentrations, and the model group and normal group were intraperitoneally injected with normal saline. The last administration was performed on the d 6 after infection. Mice inoculated with 350 TCID_50_ virus solution after administration were left untreated, and their status was observed daily and body weight was recorded in relation to death until d 14 after vaccination; mice inoculated with 25 TCID_50_ virus solution were sacrificed 3 h after administration., collect the serum, take the heart tissue and cut it into 2 parts on average, one part is homogenized in the RL1 reagent of the animal total RNA extraction kit (Chengdu Fuji Biotechnology Co., Ltd., RE-03011), and the other part is stored at -80°C for total protein extraction. The thymus and spleen were dissected, and the cardiac index, thymus index, and spleen index were calculated using the following formulas^(23)^: Organ index=Organ weight (mg) ÷Body weight (g) ×100%

### Histological staining

The mice were killed, and heart tissue was taken and fixed in 10 % formaldehyde solution. After dehydration, washing, and embedding in paraffin to prepare pathological sections, the implanted hearts were cut into 5 μm sections and stained with hematoxylin and eosin (H&E) ^(5)^.

### Cytotoxicity test

Monolayer HT-29 cells were treated with different concentrations of TSS and incubated at 37°C in a 5% CO_2_ environment for 72 h, and the cell viability was detected by CCK-8 method^(24)^. The TC_50_ (ie, the median toxicity concentration) was determined by calculating percent cell viability as a function of compound concentration using SPSS 26.0 software^(25)^.

### CPE reduction assay

To assess the way in which TSS works, we selected the intervention effect of administration at three different time points in our experiments. (1) Pre-incubation treatment: TSS was pre-added to the monolayer cells for 4 h at 37°C, followed by the addition of virus solution (100 TCID_50_) for 1h at 37°C, removal of all supernatants, and the addition of maintenance fluid. (2) Direct virucidal effect: TSS was treated with virus liquid (100 TCID_50_) at the same time, the mixture was placed in 4°C to react for 4h, and the mixture was added to the cells. After 1h, all supernatants were removed and cell maintenance fluid was added. (3) Therapeutic effect: after adsorption of CVB3 (100 TCID_50_) for 1 h, the maintenance fluid containing TSS was added to the cells. The plates were incubated at 37°C in a 5% CO_2_ incubator for 72 h. CPE was then assessed using the CCK-8 method as above. The IC_50_ was calculated from the resulting spectrophotometric data^(25)^. These experiments used were repeated at least three times.

### Viral plaque assay

The CVB3 virus solution was diluted (100 PFU) and added to a 6-well plate (6×10^5^/well). After culturing for 1 h, the cells were rinsed twice with DPBS, covered with 1.6 % low melting point agarose medium containing drug for 72 h, fixed with 4 % paraformaldehyde for 30min, and stained with crystal violet for 15 min. A detailed count of viral plaques was performed. All tests were repeated at least three times.

### Viral load measurement by qRT-PCR

The virus solution was discarded after the cells were infected with virus (100 TCID_50_) for 1 h, and different drug containing maintenance fluid were added, the test drugs plus different concentrations of TSS, the positive control group plus guanidine hydrochloride, and the normal control group with the model group were added maintenance fluid without drug. The plates were incubated at 37°C in a 5% CO_2_ incubator. Total RNA was isolated after 72 h. The primer sequences for the CVB3 viral gene were 5 ’-ccctgaatgcggctaatcc-3’ (sense) and 5 ’- attgtcaccataagcagcca-3’ (antisense), and the human GAPDH Gene was used as an internal control for cellular RNA. Amplification conditions were as follows: 95°C, 30 s, 1 cycle; 95°C, 5 s, 60°C, 30 s, 40 cycles. qRT-PCR was performed using the real-time PCR instrument Stepone with a normalized expression pattern of (2^-ΔΔCt^)。

The viral load of CVB3 infected mouse heart tissues was measured by qRT-PCR method as above. The heart tissue of mice sacrificed on the 6th day after inoculation was placed in the animal total RNA extraction kit RL1 reagent and homogenized to extract RNA. Primer sequences for the CVB3 viral gene are as above, and the mouse GAPDH Gene was used as an internal reference for tissue RNA. The amplification conditions are as above.

### TLR3 and its downstream signaling factors expression detection by qRT-PCR

Preparation of cell samples and mouse heart tissue samples was as above. Sequences are given in table1 and table2. The human and mouse GAPDH genes, respectively, were used as internal references. The amplification conditions are as above. qRT-PCR was performed using the real-time PCR instrument Stepone with a normalized expression pattern of (2^-ΔΔCt^)。

**Table 1.**
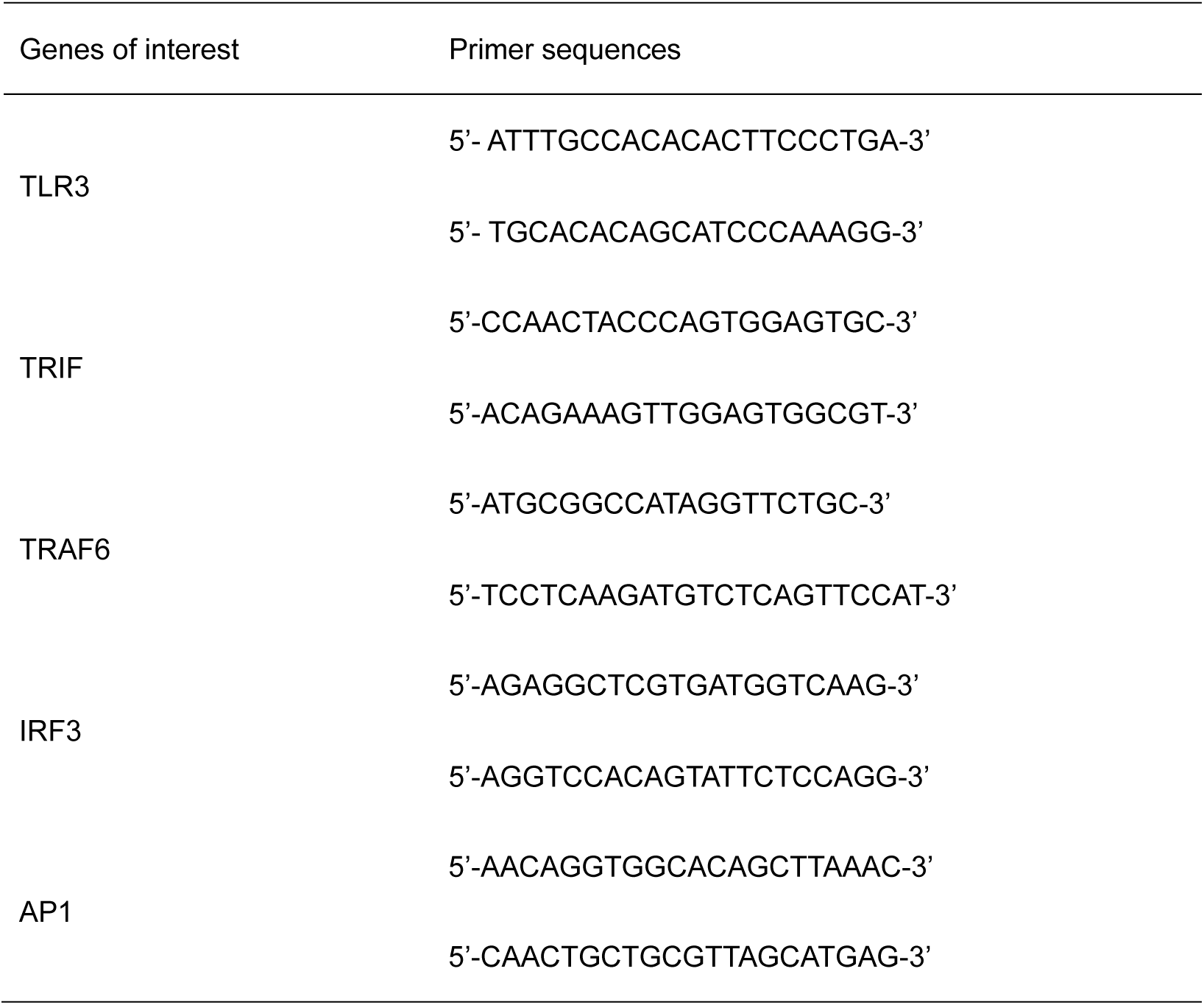
Cell Primer sequences

**Table 2.**
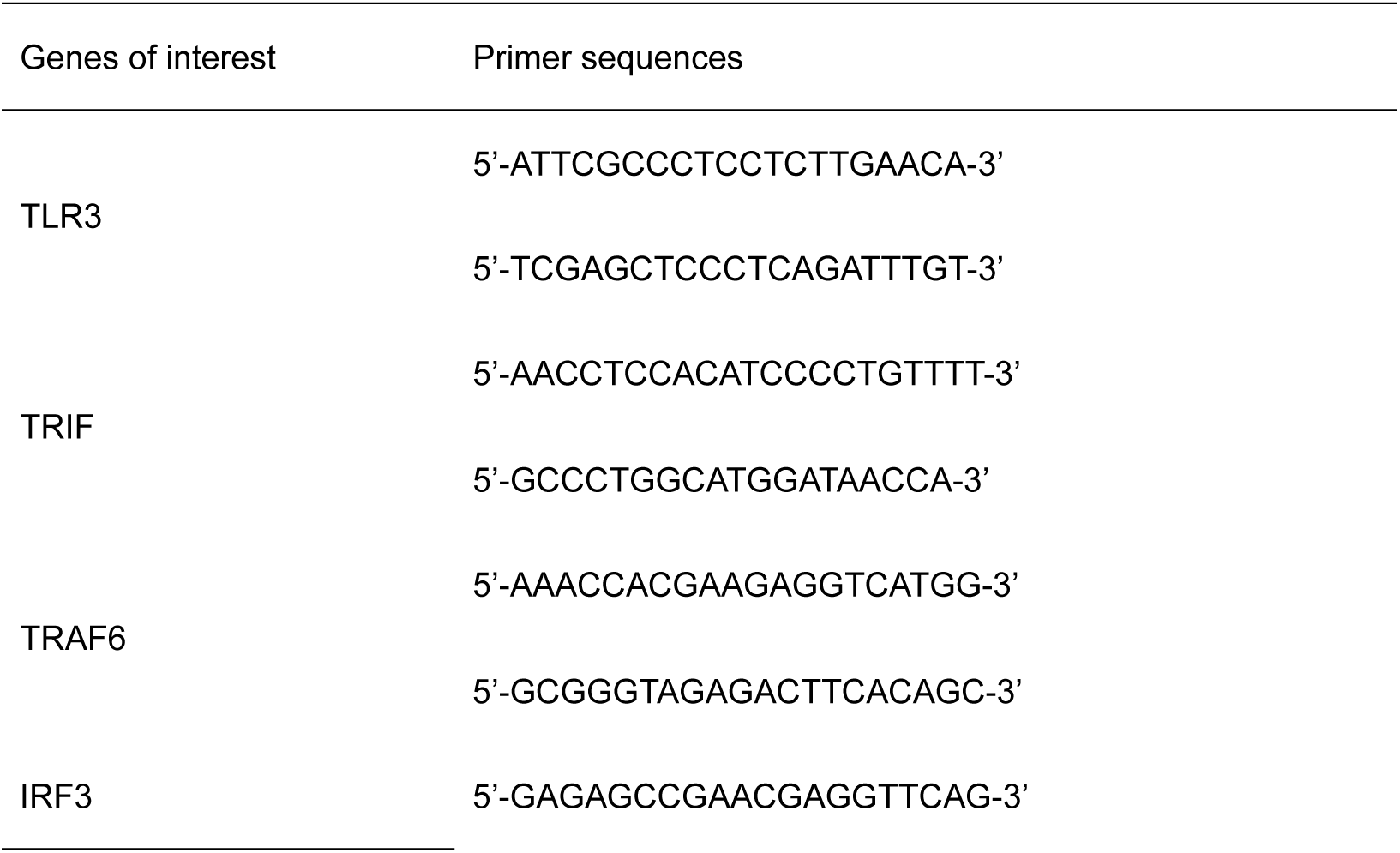
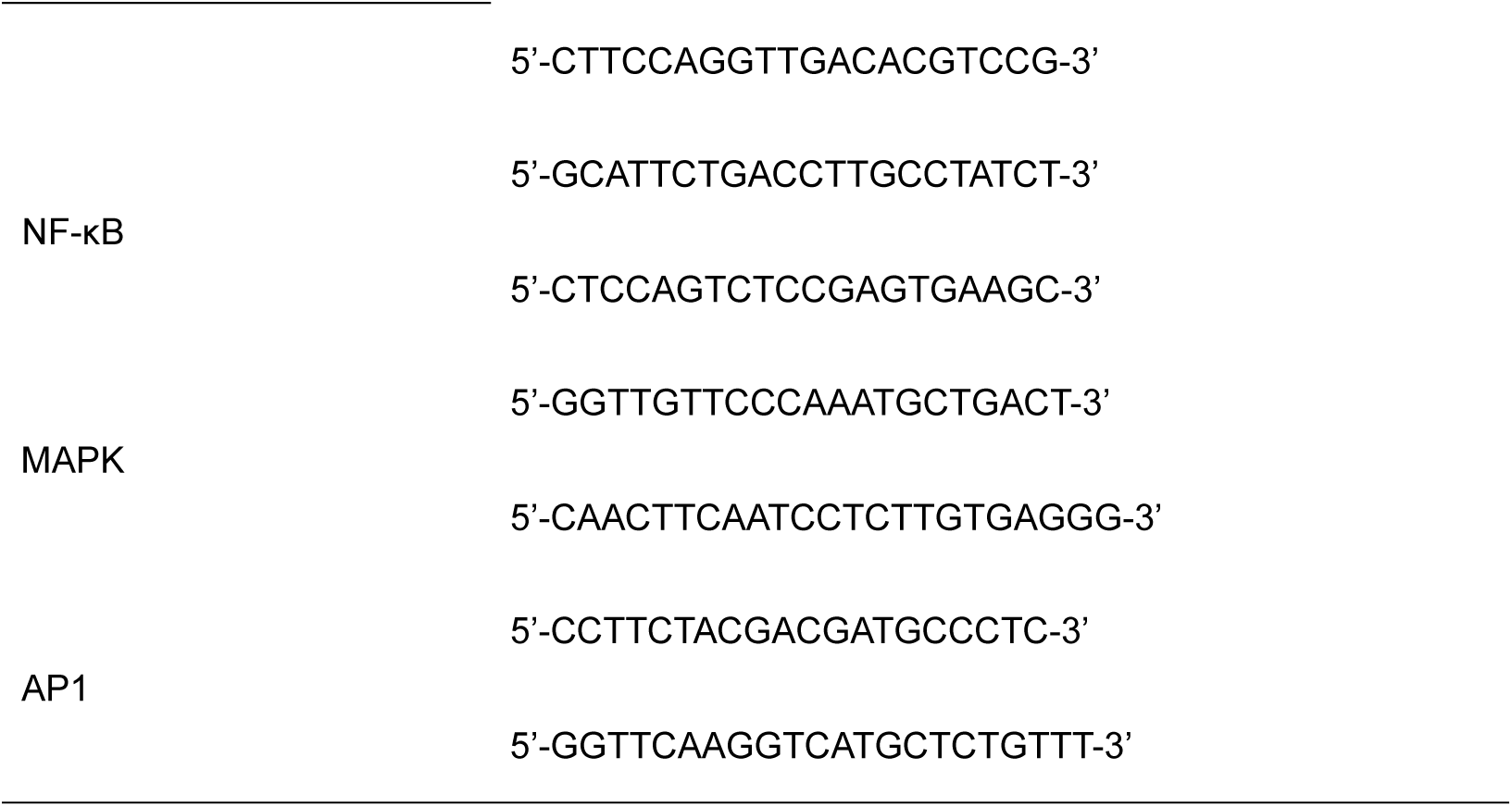
Mouse primer sequence

**Table 3.**
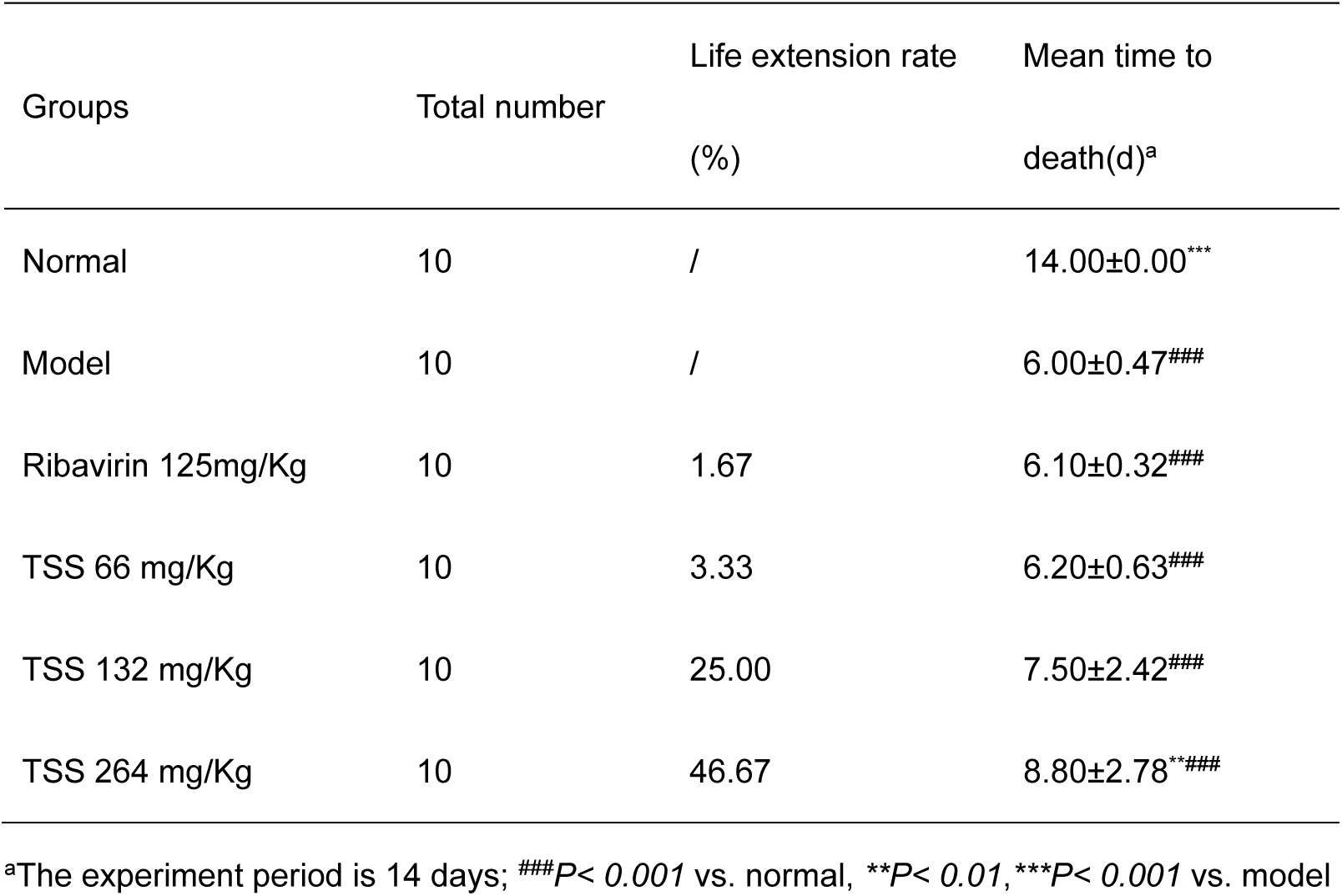
Protective effect of TSS on CVB3 virus-infected mice (n=10)

### Cytokine secretion assay (ELISA)

After cells were infected with virus (100 TCID_50_) for 1 h, different drug containing maintenance fluid were added. The plates were incubated at 37°C in a 5% CO_2_ incubator for 72 h. Collect cell supernatant. The expression levels of NF-κB, MAPK, IL-6, TNF-α and IFN-β in cell supernatants were measured by ELISA kits (Nanjing Jiancheng Bioengineering Institute, China). Mouse serum was collected on the d 6 after virus inoculation. The expression levels of IL-6, TNF-α and IFN-β in mouse serum were detected by ELISA kit.

### Western Blot (WB)

The expressions of TLR3,TRIF in heart tissues of CVB3 infected mice were measured by Western blot. Proteins were extracted from heart tissues of mice sacrificed at d 6 after inoculation. Mouse GAPDH was used as an internal reference for tissue proteins.

### Statistical analysis

SPSS statistics 26.0 software was used to analyze the data, one way ANOVA was performed to analyze the data between groups, and *P < 0.05* was considered statistically significant, and Graphpad Prism 8.0 (Graphpad Software Inc, USA) software was used to draw images.

## RESULTS

### Therapeutic effect of TSS on CVB3 virus in vivo

To verify the pharmacological effects of TSS, we evaluated the therapeutic effect of TSS on CVB3-induced myocarditis in mice. As shown in Figure 3.a, the life extension rate of mice injected with TSS 66 mg/Kg, 132 mg/Kg and 264 mg/Kg by sc injection was 46.67 %,25.00 % and 3.33 %, respectively, while the mice injected with normal saline all died within 6d. The positive control drug ribavirin had limited protective effect, and the life extension rate of sc injection of ribavirin was 1.67 %. Furthermore, TSS was able to reduce weight loss (Fig. 3b). The mean time to death of mice is shown in Table 1. Compared with the model group (mean time to death 6.00±0.47 d), the mean time to death (8.80±2.78 d) of TSS 264 mg·Kg^-1^·d^-1^ was significantly prolonged (*P<0.01*). Ribavirin prolongs the mean time to death of mice to 6.10±0.32 d.

**Fig. 3.**
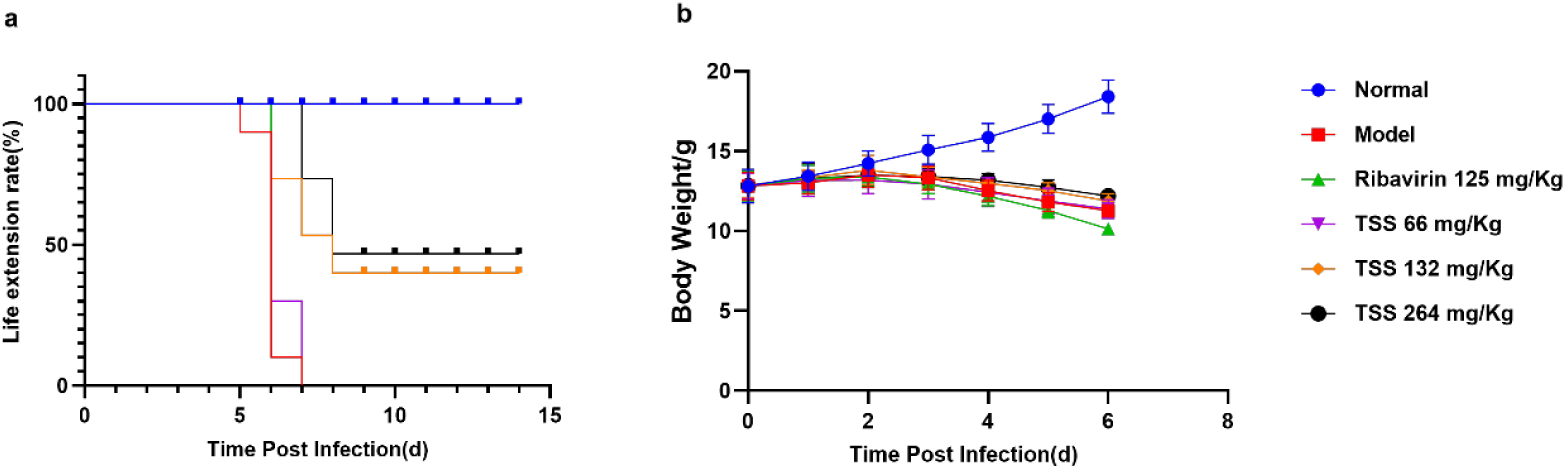
**a** The life extension rate: mice were continuously observed for 14 days since virus inoculation. **b** Body weight of mice (n=10 each group).

### TSS inhibits CVB3 viral replication and protects mouse heart

In this study, qRT-PCR was used to detect the expression level of viral mRNA in the heart tissue of CVB3 virus-infected mice. Using mouse GAPDH as an internal reference gene, relative quantification by qRT-PCR was 2^-ΔΔCt^ method was used to compare the expression results. As shown in Fig. 4a, the CVB3 gene expression level decreased in all drug-treated groups. This means that all drugs can effectively inhibit CVB3 virus replication in mice with viral myocarditis. Compared with the model group, TSS 264mg/Kg (*P < 0.001*), 132mg/Kg (*P < 0.001*) and 66mg/Kg (*P<0.001*) could significantly down-regulate the expression of CVB3 mRNA.

**Fig. 4.**
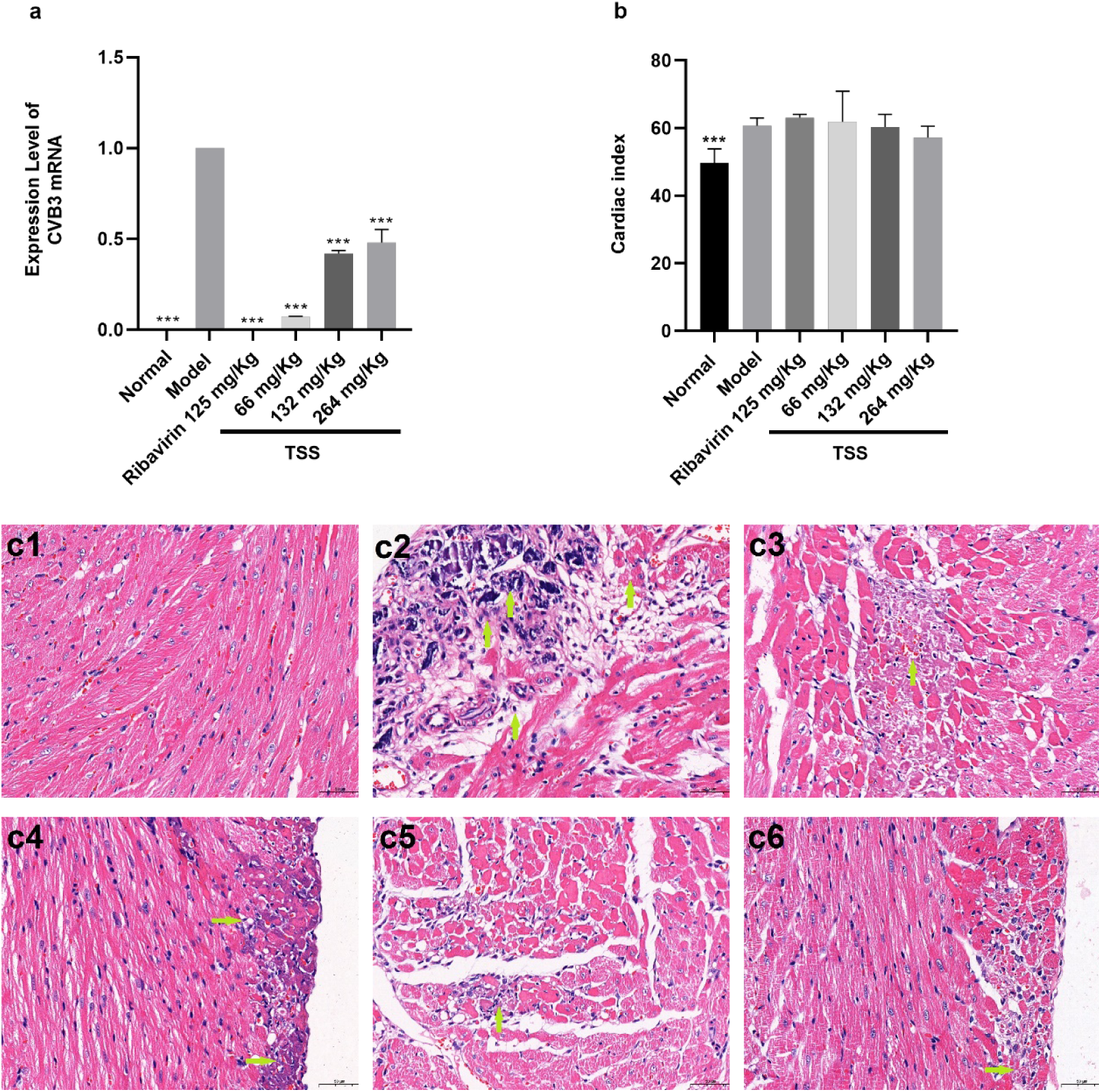
The protective effect of drugs on the cardiac tissue of CVB3 virus-infected mice.**a** CVB3 mRNA levels in mouse hearts measured by qRT-PCR and normalized to GAPDH. TSS treatment can decrease the expression of CVB3 mRNA(*\*\*\*P< 0.001* vs. model). **b** Cardiac index. *\*\*\*P< 0.001* vs. model. **c** H&E staining of sectioned hearts (×400). Pathological changes were evaluated according to the structure of cardiac endothelial cells, myocardial fiber necrosis, calcium salt precipitation, and inflammatory cell infiltration. **c1** normal group: the structure of epicardium and endocardial endothelial cells is complete and clear, the shape of myocardial fibers is relatively normal, no obvious degeneration and necrosis; **c2** model group: local myocardial fibrosis and necrosis, epicardium and media layer myocardial fiber foci The necrosis and calcification were more obvious in the local area, and purple-black calcium salt deposition, a small amount of fibrous tissue and lymphocytes were seen in the necrotic area; **c3** Ribavirin control group: myocarditis lesions were significantly alleviated, and only a small amount of myocardial fiber degeneration and necrosis; **c4** TSS 66 mg·Kg^-1^·d^-1^; **c5**TSS 132 mg·Kg^-1^·d^-1^; **c6** TSS 264 mg·Kg^-1^·d^-1^; (TSS can significantly reduce the inflammatory lesions of myocardial tissue, and a small amount of myocardial fibrosis and necrosis)

As shown in Fig.4b, the cardiac index data showed that TSS 264mg/Kg had a significant protective effect on viral myocarditis on the d 6 after infection (*P<0.01*). Histopathological examination (H&E staining) further showed that TSS significantly reduced the inflammation in mouse heart tissue, as shown in Fig. 4c.

mRNA levels in mouse hearts measured by qRT-PCR and normalized to GAPDH. TSS treatment can decrease the expression of CVB3 mRNA(*\*\*\*P< 0.001* vs. model). **b** Cardiac index. *\*\*\*P< 0.001* vs. model. **c** H&E staining of sectioned hearts (×400). Pathological changes were evaluated according to the structure of cardiac endothelial cells, myocardial fiber necrosis, calcium salt precipitation, and inflammatory cell infiltration. **c1** normal group: the structure of epicardium and endocardial endothelial cells is complete and clear, the shape of myocardial fibers is relatively normal, no obvious degeneration and necrosis; **c2** model group: local myocardial fibrosis and necrosis, epicardium and media layer myocardial fiber foci The necrosis and calcification were more obvious in the local area, and purple-black calcium salt deposition, a small amount of fibrous tissue and lymphocytes were seen in the necrotic area; **c3** Ribavirin control group: myocarditis lesions were significantly alleviated, and only a small amount of myocardial fiber degeneration and necrosis; **c4** TSS 66 mg·Kg^-1^·d^-1^; **c5**TSS 132 mg·Kg^-1^·d^-1^; **c6** TSS 264 mg·Kg^-1^·d^-1^; (TSS can significantly reduce the inflammatory lesions of myocardial tissue, and a small amount of myocardial fibrosis and necrosis)

### TSS inhibits the expression of inflammatory markers in mice

LDH, CK-MB and cTn-Ⅰ are three enzymes, which are commonly used in the detection of myocardial inflammation or injury. As shown in Fig. 5a, Coxsackie virus infection caused an increase in the expression levels of LDH, CK-MB, and cTn-I, and all drug groups showed different degrees of myocardial enzyme inhibition (*P<0.05*). These data suggest that irisin sodium sulfonate can protect myocardial injury in mice with viral myocarditis.

**Fig. 5.**
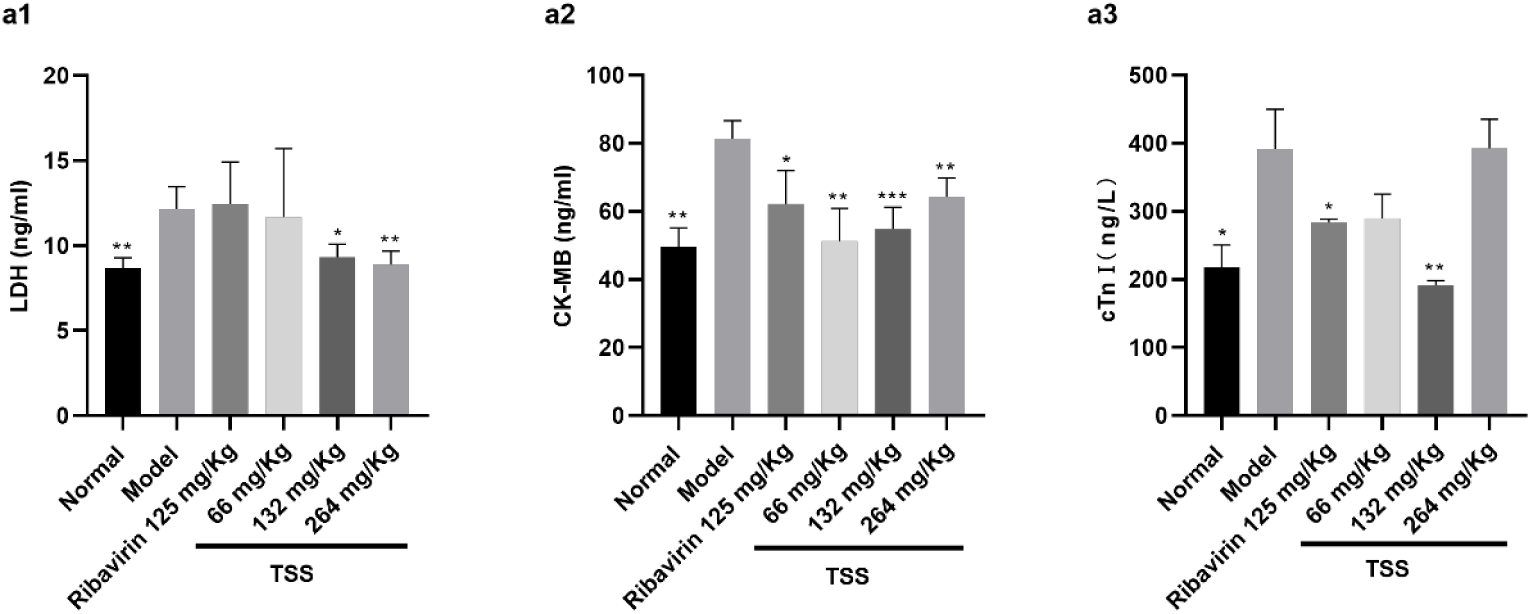
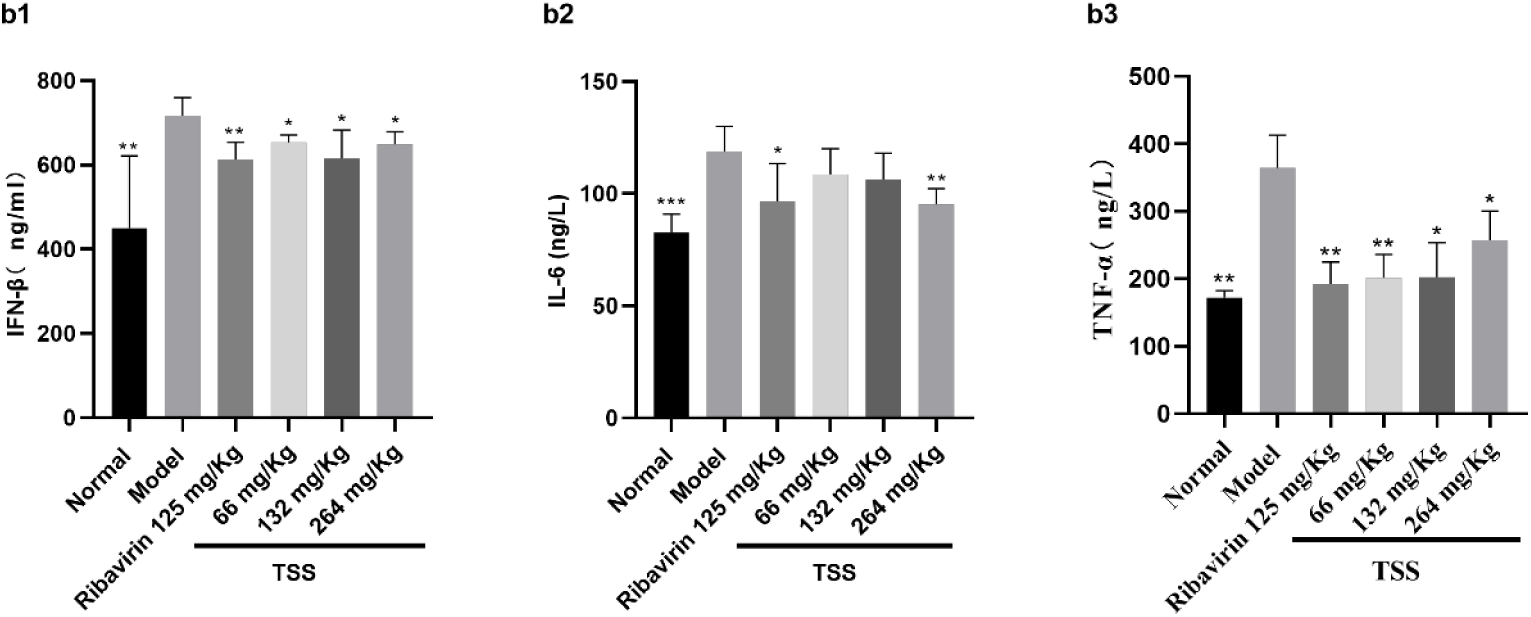
The expression levels of myocardial enzymes and inflammatory factors in the heart tissue of infected mice were detected. Heart tissue homogenate was prepared, and enzyme-linked immunosorbent assay (ELISA) was used to detect myocardial enzymes LDH, CK-MB, cTnⅠ and inflammatory cytokines IFN-β, IL-6 and TNF-α. *\*P< 0.05, **P< 0.01, ***P< 0.001* vs model, each group n=6

Inflammatory factors play an important role in innate and acquired immunity caused by infection. In this paper, IFN-β, IL-6 and TNF-α were detected as inflammatory markers after CVB3 infection. As shown in Fig. 5b, its expression level increased in the hearts of virus-infected mice, and the drug treatment group showed varying degrees of decreased expression levels of IFN-β, IL-6, and TNF-α (*P < 0.05*). These data suggest that TSS can modulate the inflammatory response in myocarditis.

### TSS improves immune function in mice

Mice were killed on d 6 post-infection, and spleen and thymus specimens were collected. As shown in Figure 6a, the spleen of the model group was greatly changed after infection with CVB3, and TSS was able to improve the appearance of the spleen. The spleen and thymus indexes of mice in the model group were significantly lower than those in the normal group *(P<0.01*). However, TSS (264mg/Kg) significantly improved the above indicators (*\*\*\*P<0.001, **P<0.01*), indicating that TSS can effectively inhibit the damage of immune organs.

**Fig. 6.**
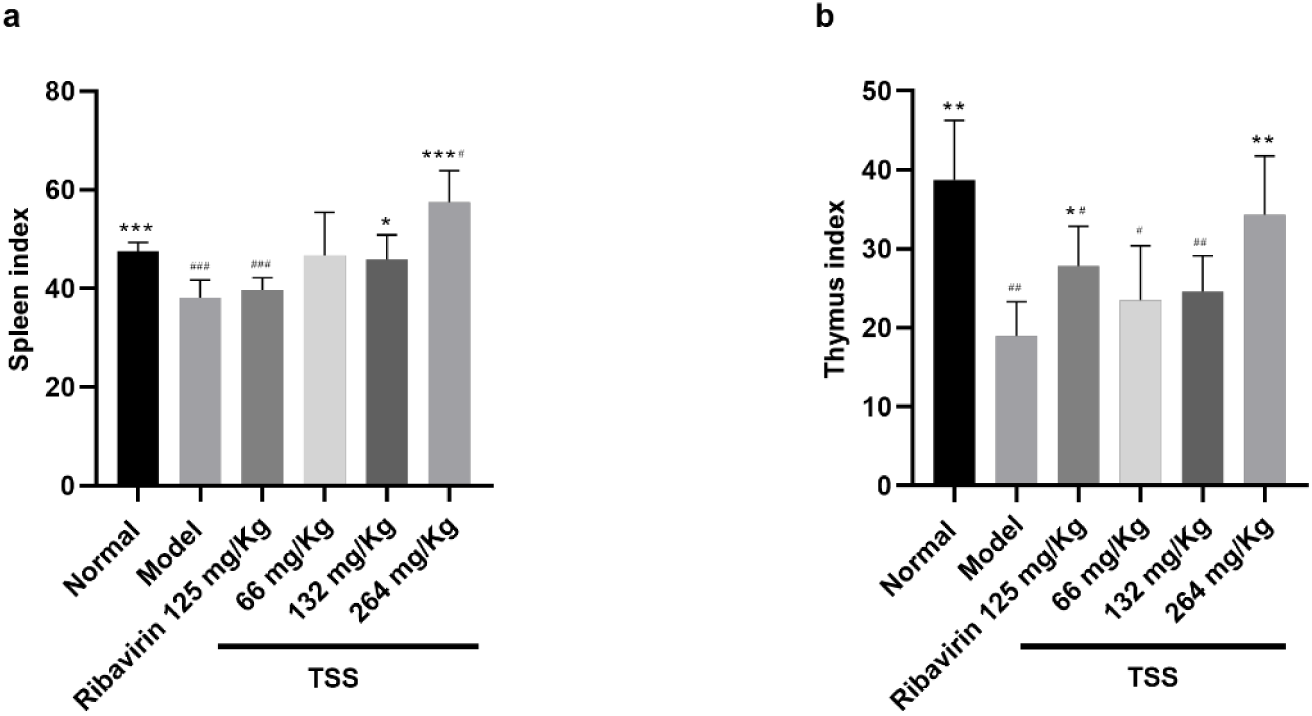
**a** Spleen index of infected mice after 6 days. **b** Thymus index. *\*P< 0.05, **P< 0.01, ***P< 0.001* vs model., *^#^P< 0.05,^##^P< 0.01,^###^P< 0.001* vs normal, each group n=6

### Effects of TSS on the TLR3 pathway

TLR3 plays an important role in virus recognition and innate immunity. It can recognize the viral intermediate dsRNA, and then activate the expression of its downstream immune-phase pathway-related factors. In this paper, TLR3 and its downstream TRIF, TRAF6, IRF-3, NF-κB, MAPK, AP1 were used as immune system markers to regulate virus infection. Detected that TLR3 and its downstream pathway cytokine expression increased in mice heart tissue after its downstream cytokine mRNA levels in each group were detected by qRT-PCR at d 6 after viral infection. The drug treated group showed different degrees of inhibition (*P < 0.05*) of this pathway. These data demonstrate that TSS can modulate the immune system in vivo and exert therapeutic effects through the TLR3 pathway (Fig 7).

**Fig. 7.**
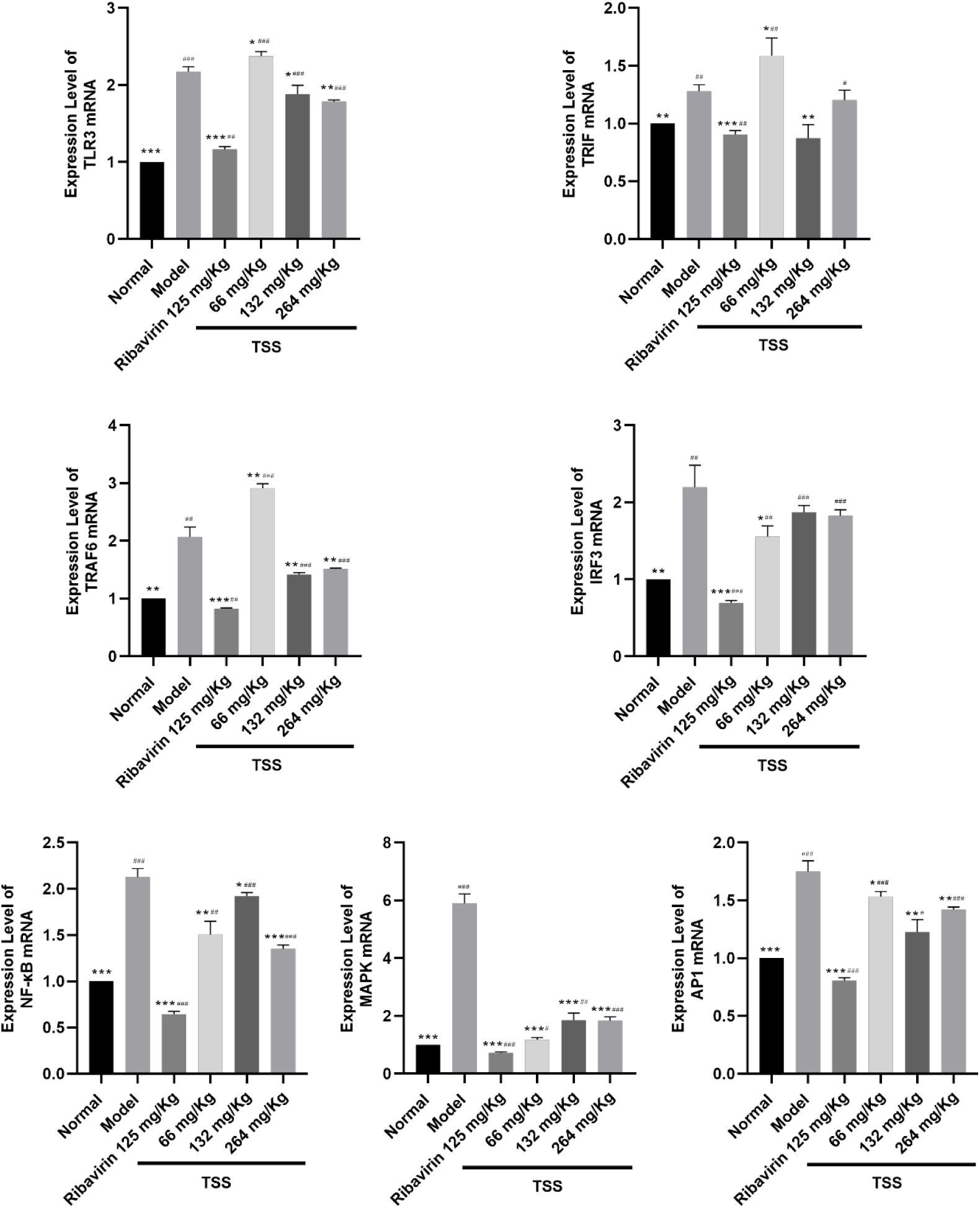
TSS reduces the expression levels of TLR3, TRIF, TRAF6, IRF-3, NF-κB, MAPK, AP1 mRNA in the heart tissue of CVB3-infected Balb/c mice. TSS was added to 25 TCID50 CVB3 viral fluids to infect animals at a series of concentrations of 66, 132 and 264 mg/Kg. TLR3 and its downstream cytokine mRNA levels in each group were detected by qRT-PCR at d 6 after viral infection. *\*P< 0.05, **P< 0.01, ***P< 0.001* vs model; *^#^P< 0.05,^##^P< 0.01,^###^P< 0.001* vs normal. In total, three independent experiments were performed.

**Fig. 8.**
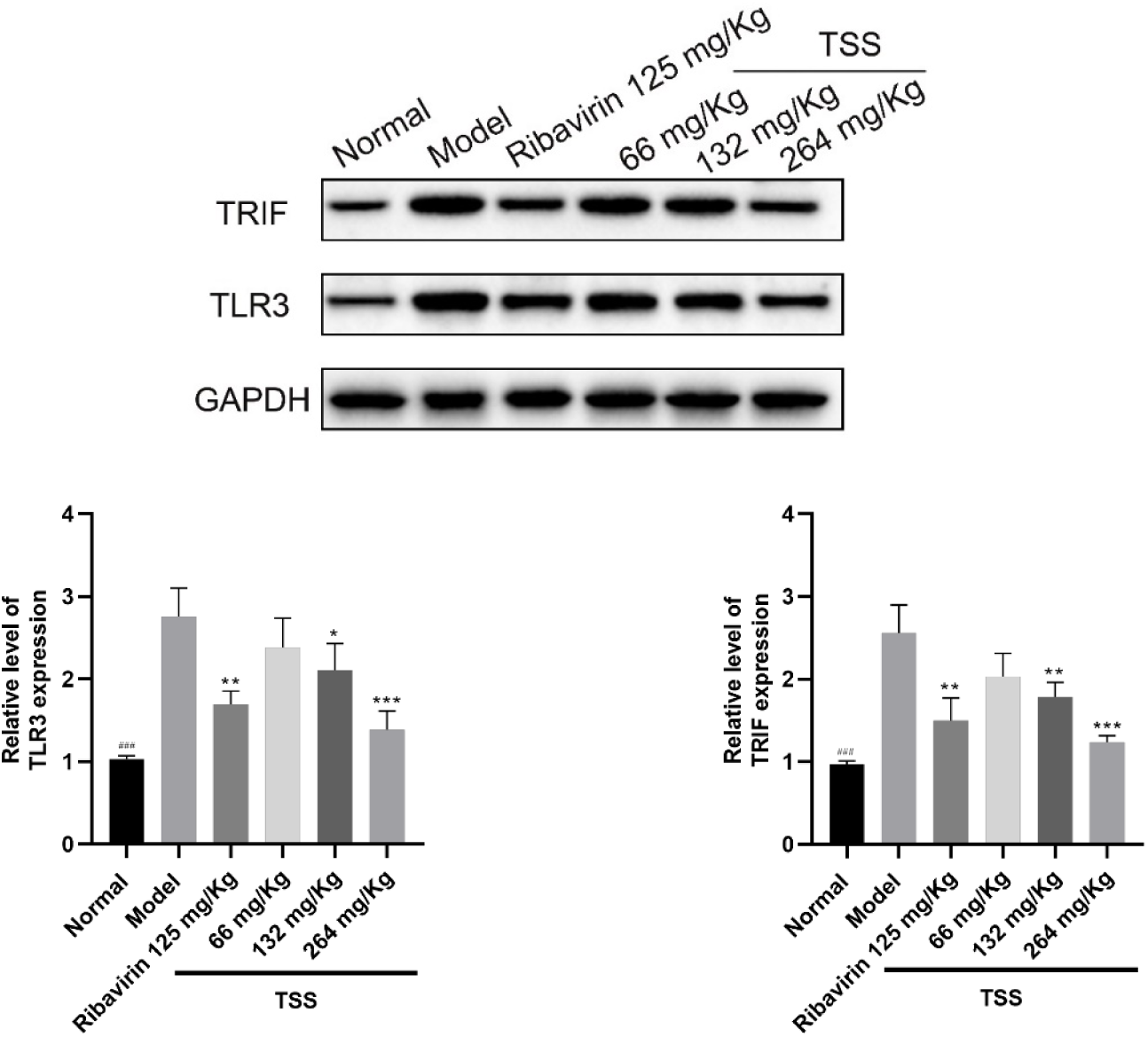
In-depth analysis of the effect of TSS inhibition of TLR3 pathway on CVB3 infection. Mice were infected with CVB3 virus and treated with TSS to observe the effect of TSS on TLR3 and TRIF protein levels. *\*P< 0.05, **P< 0.01, ***P< 0.001* vs model; *^###^P< 0.001* vs model.

### In vitro antiviral activity of TSS on CVB3

The cytotoxicity of TSS on HT-29 cells was detected by CCK-8 assay. The results showed that TSS had no obvious cytotoxic effect on HT-29 cells, with a TC_50_ value of 1.5309 mM. To verify the in vitro anti-CVB3 viral activity of TSS, the inhibitory effect of TSS on CVB3 induced cytopathic effect (CPE) was evaluated on HT-29 cells. The stage at which TSS exerted antiviral effects was determined by different modes of administration. TSS were added to HT-29 cells at three different time points: Drug pre-incubation for 4h (preventive effect), simultaneous treatment of drug and virus (direct killing effect of drug on virus) and after virus entry (therapeutic effect). As shown in figure 9a-c, TSS exerted an antiviral effect by entering cells, with an IC_50_ value of 13.8529 μM. Its concentration was positively correlated with the corresponding inhibitory effect.

**Fig. 9.**
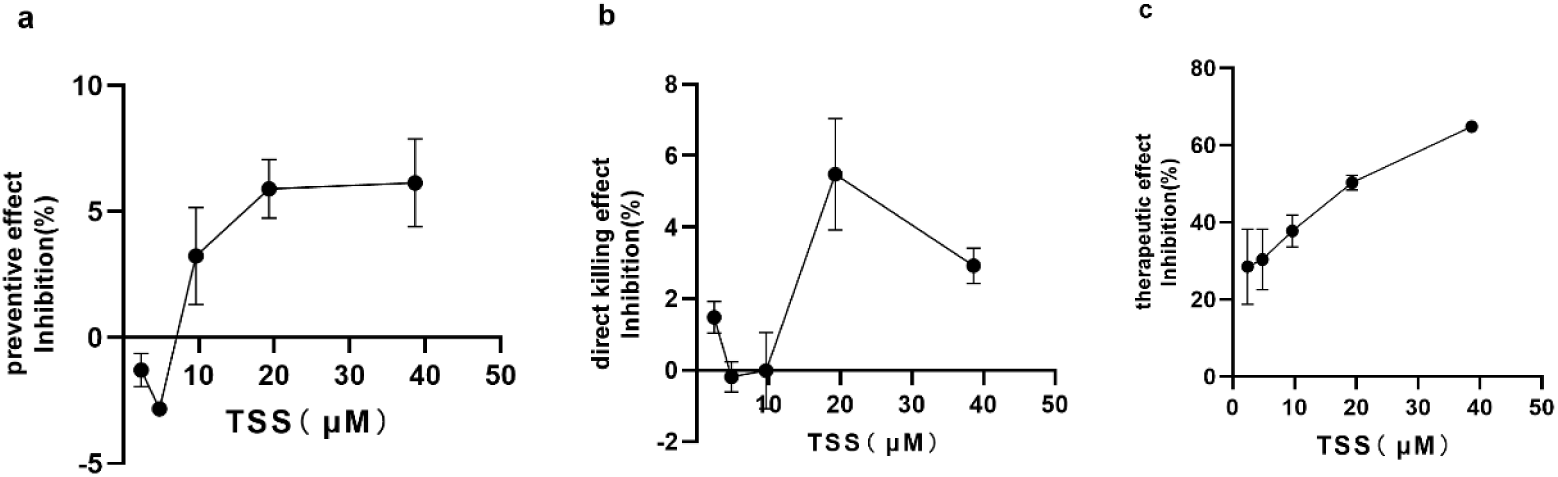
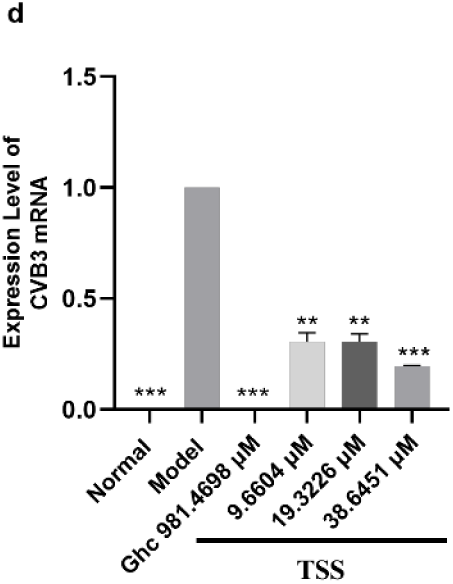
Antiviral activity of TSS on CVB3 viruses in vitro. **a-c** Inhibition rates of TSS (from 2.4125 μM to 38.6451 μM) against CVB3. CVB3 with different infection protocols in HT-29 cells. Each concentration took up three wells for each assay, and three independent determinations were carried out, determined by CCK-8 assay. **d** CVB3 mRNA levels in HT-29 cells measured by qRT-PCR and normalized to GAPDH. TSS treatment can decrease the expression of CVB3 mRNA(*\*\*\*P< 0.001* vs. model).

**Fig. 10.**
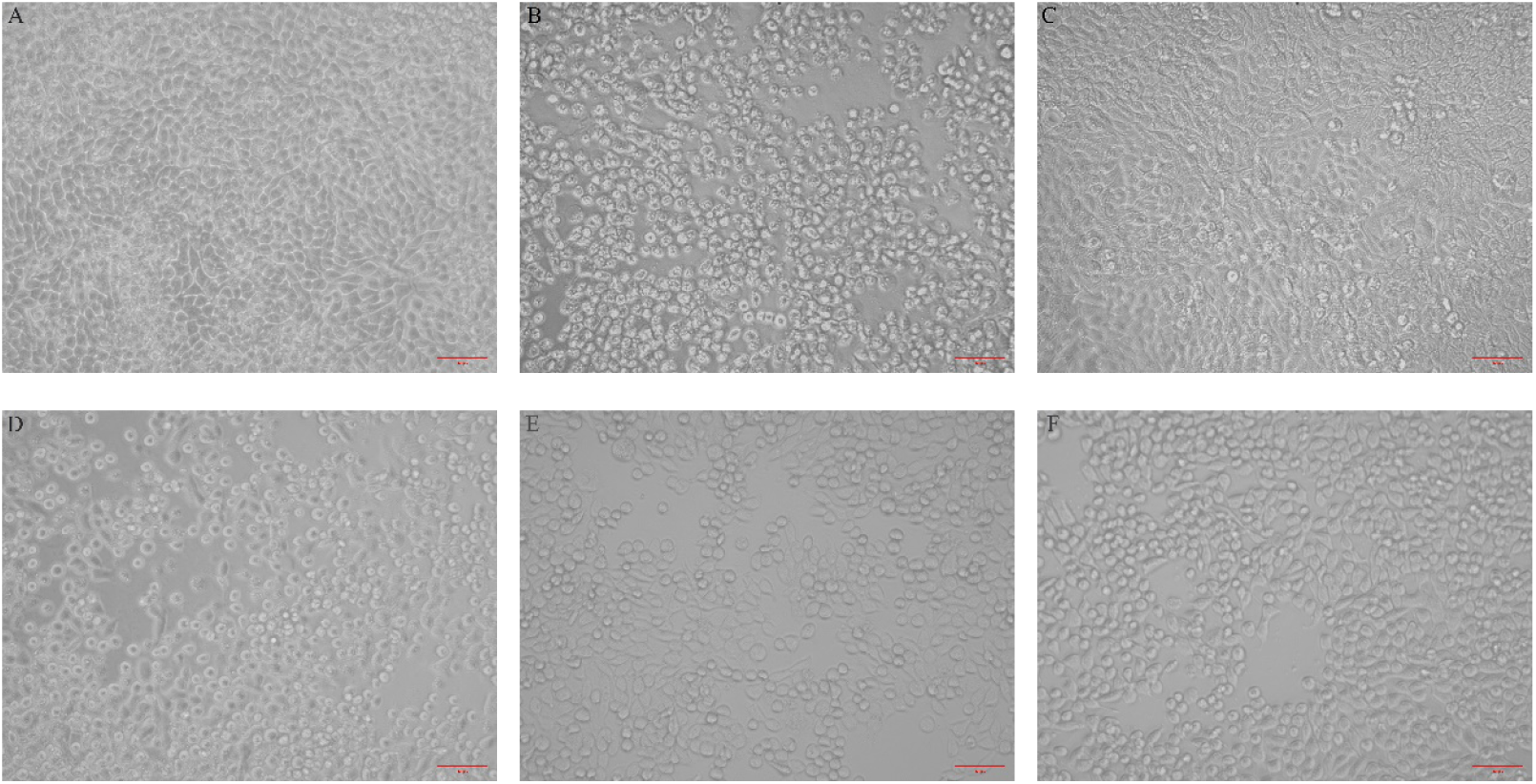
Effect of TSS on the morphology of CVB3 infected HT-29 cells (therapeutic effect). After the cells were inoculated with CVB3 virus for 72 h, the normal group could be observed under the microscope (**a**) the cells were adherent and in good growth status, did not deformation, arranged closely, did not undergo shedding; Model group (**b**) cells shrunk and a large number of cells were detached, and the remaining adherent cells refractoriness was high; After treatment with the positive drug Ghc (**c**) cytopathic changes decreased, there were a few cells refractoriness increased; The groups treated with different concentrations showed different degrees of lesions in a dose-dependent manner, most cells were adherent and morphologically normal after treatment with high dose (**f**) of TSS, and some cells were detached, while the lesions of cells treated with medium dose (**e**) versus low dose (**d**) in the higher dose group were severe, and a large number of cells shrunk and sloughed off, producing cell debris.

**Fig. 11.**
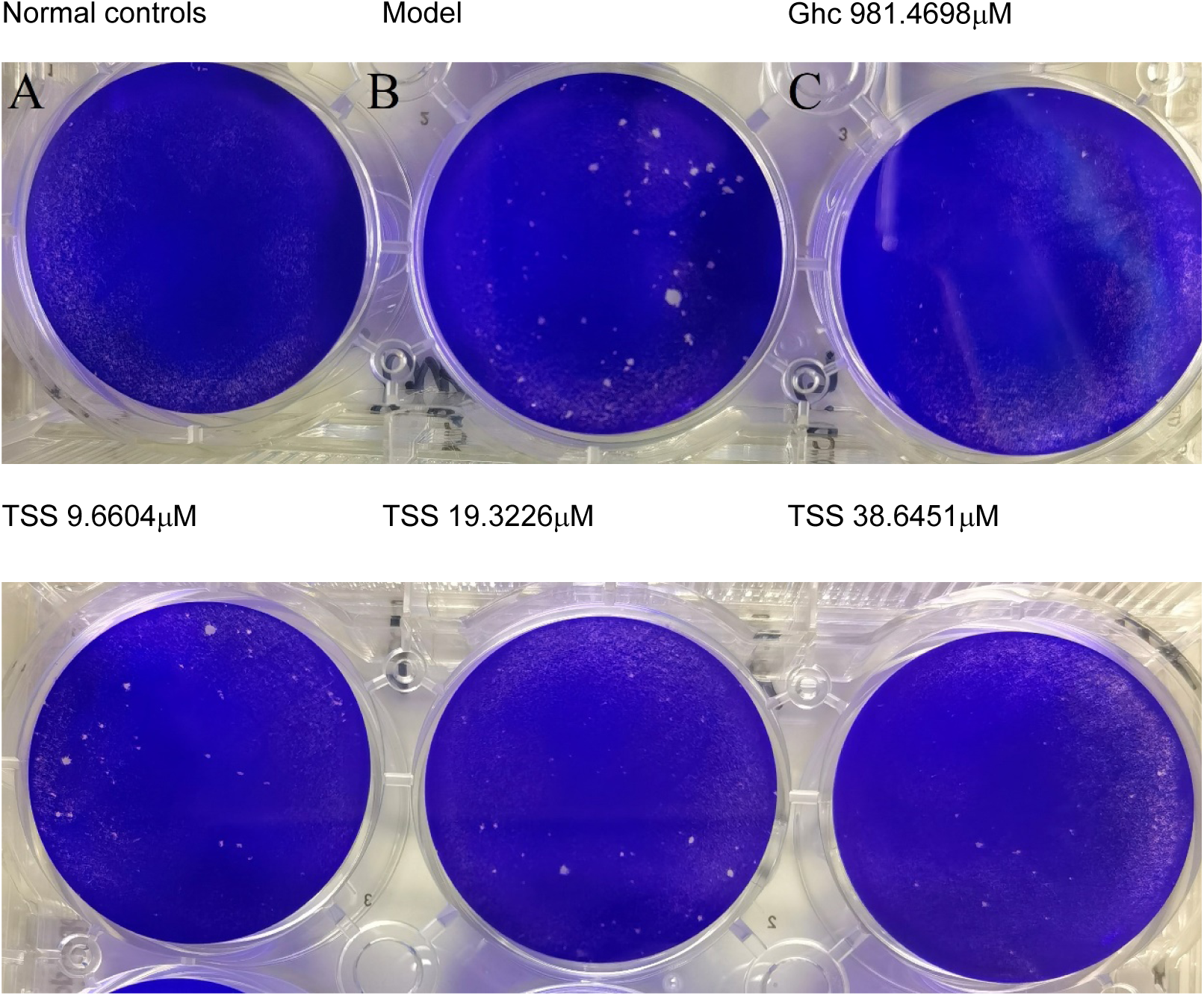
The effect of TSS on plaque formation in HT-29 after CVB3 infection. Cells in the blank control group had normal morphology and did not form empty spots; The model group developed lesions forming about 50 plaque units; Plaque formation was significantly reduced after guanidine hydrochloride treatment (*P < 0.05*), and the plaque number was similarly significantly reduced after TSS treatment (*P < 0.05*) in a dose-dependent manner, with plaque number TSS 38.6451 μM < TSS 19.3226 μM < TSS 9.6604 μM).

To investigate whether TSS affected the replication of virus in host cells, HT-29 cells were infected with CVB3 virus with or without TSS treatment. The concentrations of TSS in the treatment groups were 9.6604 μM, 19.3226 μM, and 38.6451 μM. 72 h after infection, the expression of viral genes was detected by qRT-PCR using primers against CVB3 mRNA. As shown in Figure.9d, total RNA was isolated from infected cells, and CVB3 mRNA expression was highest in the model group cells without TSS treatment, and TSS significantly reduced its expression in a dose-dependent manner. There was no CVB3 mRNA expression in the normal group (*P < 0.001*). Compared with the model group, the mRNA expression levels of CVB3 in the TSS drug group at the doses of 9.6604 μM, 19.3226 μM and 38.6451 μM were significantly decreased (*P < 0.001*).

### TSS inhibits the expression of inflammatory markers in HT-29 cells

Infected HT-29 cells with CVB3, the expression levels of inflammatory markers in the cells increased. As shown in Figure.12, the expression levels of IFN-β, IL-6 and TNF-α in the drug treatment group were improved to different degrees (*P<0.05*). These data suggest that TSS can modulate the inflammatory response of CVB3-infected cells.

**Fig. 12.**
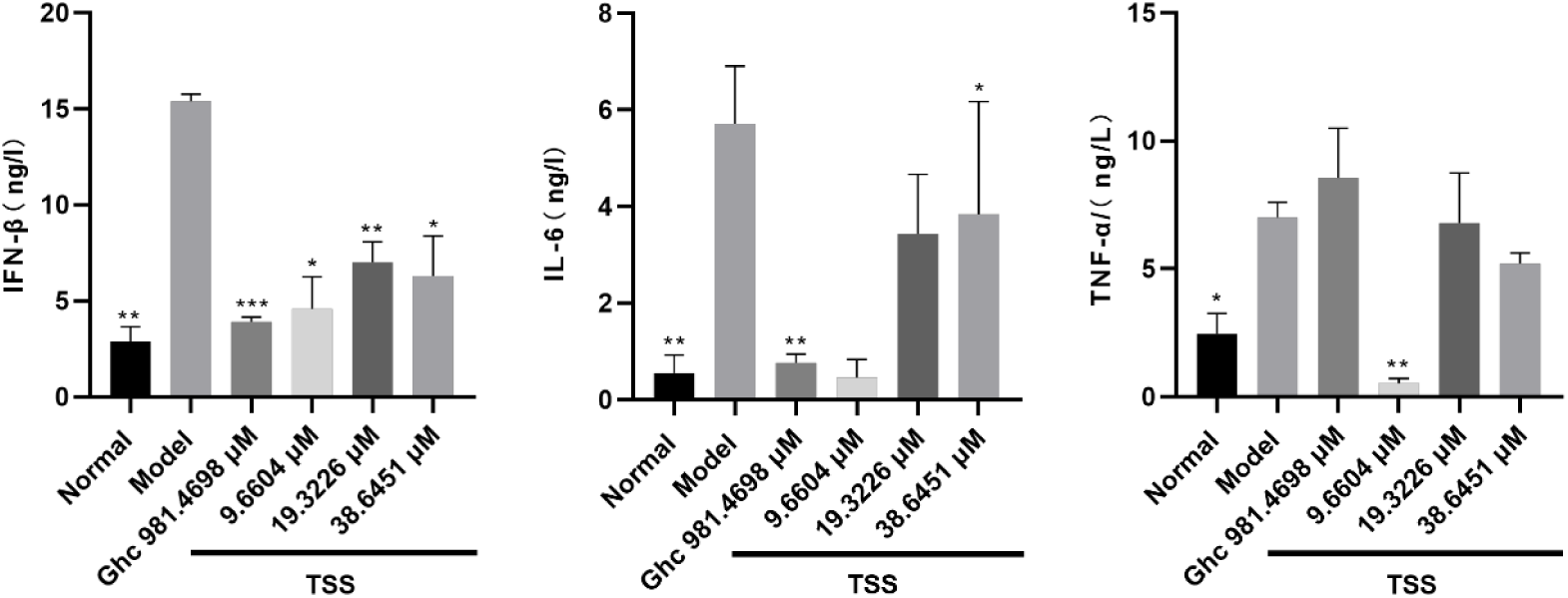
Detection of the expression levels of inflammatory factors in infected HT-29 cells. The cell supernatant was prepared for detection of inflammatory cytokine IFN-β、IL-6、TNF-α by ELISA. *\*P< 0.05, **P< 0.01, ***P< 0.001* vs model, each group n=3

### Effects of TSS on the TLR3 pathway

In virus-infected cells, the expression of TLR3 and its downstream cytokines in cells increased, and the drug treatment group showed different degrees of inhibition of this pathway (*P<0.05*). These data suggest that TSS can exert antiviral effects through the TLR3 pathway (Fig.13).

**Fig. 13.**
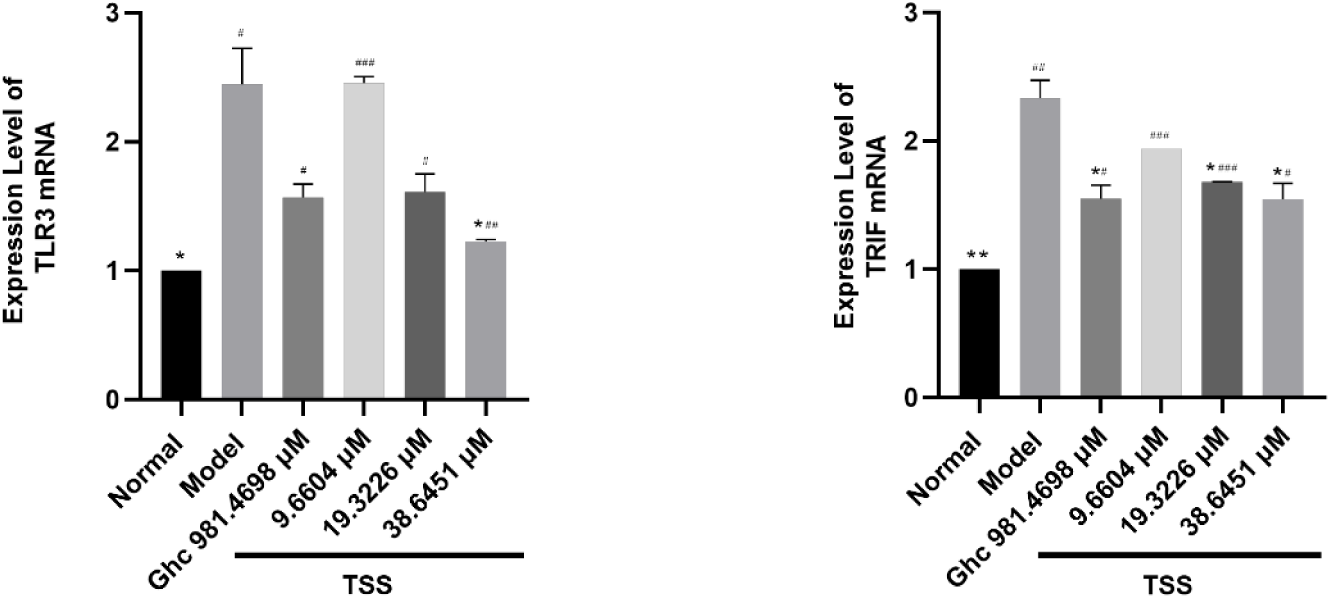
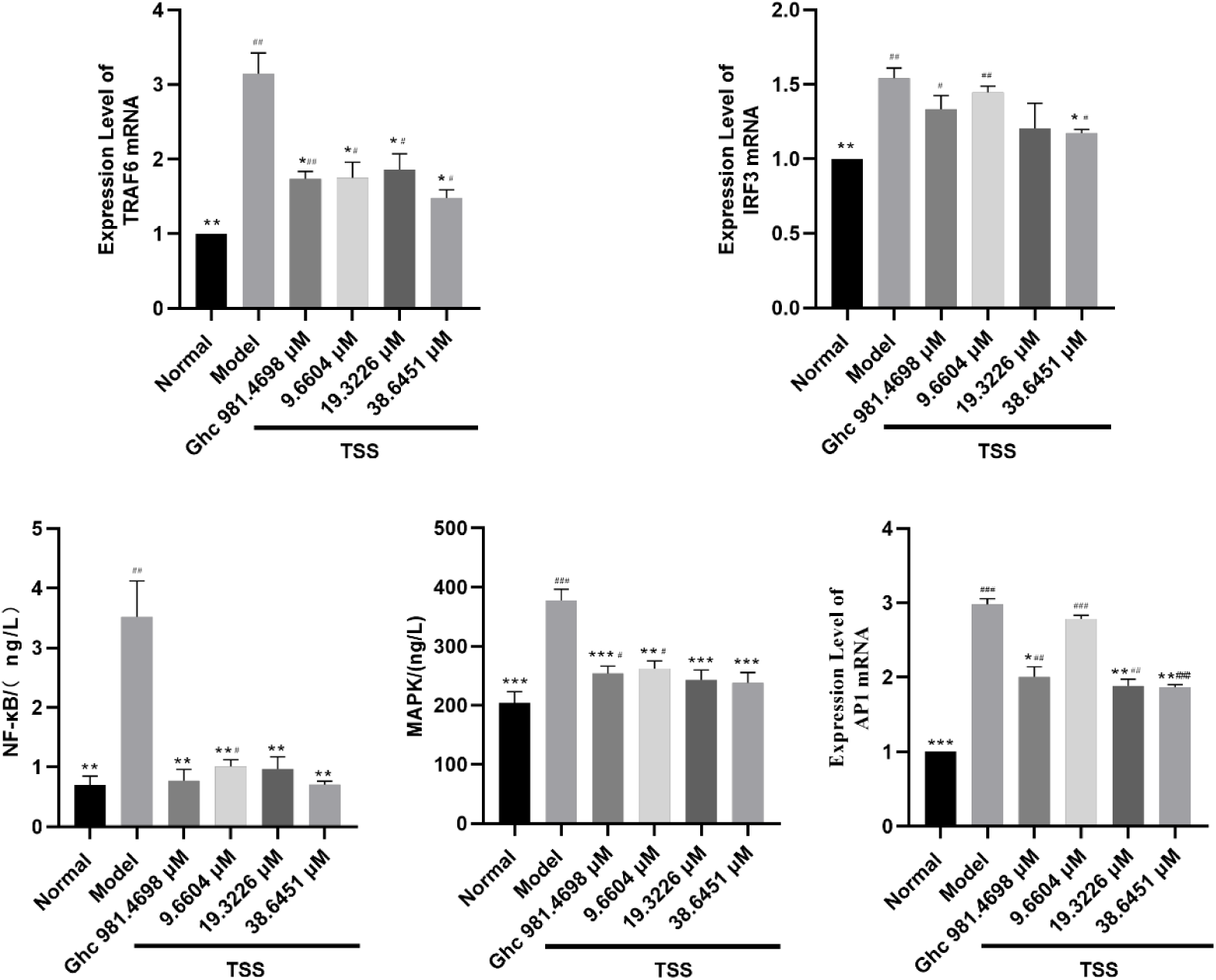
TSS reduces the expression levels of TLR3, TRIF, TRAF6, IRF-3, NF-κB, MAPK, AP1 in CVB3-infected HT-29 cells. TSS was added to 100 TCID50 CVB3 virus fluids to infect cells at a range of concentrations of 9.6604μM, 19.3226μM and 38.6451μM. The expression levels of TLR3 and its downstream cytokines in each group were detected by qRT-PCR and ELISA at 72h after virus infection. *\*P< 0.05, **P< 0.01, ***P< 0.001* vs model; *^#^P< 0.05,^##^P< 0.01,^###^P< 0.001* vs normal. In total, three independent experiments were performed.

## DISCUSSION

Viral myocarditis (VMC) is a common clinical disease, usually caused by virus invasion of cells, with a high incidence, especially in children and adolescents. Numerous studies have shown that viruses such as enteroviruses and adenoviruses can infect the myocardium and cause symptoms of myocarditis^(26)^. Coxsackie B group is the main pathogen of viral myocarditis, of which Coxsackie virus B3 (CVB3) is the most pathogenic^(27)^. However, the infection mechanism of CVB3 is still unclear, which seriously hinders the development of effective clinical treatment. Therefore, the infection mechanism of CVB3 and the development of antiviral drugs are imminent. Traditional Chinese medicine may be a potential alternative medicine for the treatment of the disease. Recently, some studies of traditional Chinese medicine for the treatment of CVB3 infection have been carried out^(6, 28, 29)^. Tectorigenin sodium sulfonate (TSS) is a modified product of chemical synthesis of irisin from the dried rhizome of Chinese herb *Iris tectorum Maxim*, which is widely used as a folk medicine in our country to treat upper respiratory tract infections, and it is clinically used for anti-inflammatory and antiviral applications. In this study, we analyzed its antiviral effects in vivo and in vitro, and revealed and discussed its antiviral mechanism.

We examined the in vivo inhibitory activity of TSS against CVB3 virus. Sc injection of TSS has obvious inhibitory effects on CVB3-infected mice, including improving the life extension rate of mice, prolonging the mean time to death, reducing cardiac index and cardiac disease. It also has a certain effect on the secretion of inflammatory factors. It has been reported that the pathological damage caused by CVB3 infection is mainly due to the inflammatory response of the host to the virus^(3)^, and some patients who cleared the virus from the myocardium recovered completely without apparent sequelae; For others, an inappropriate immune response ensues, leading to chronic, irreversible cardiomyopathy^(30)^. In this study, H&E staining showed severe cardiac inflammation in the model group mice, including myocardial fibrosis and necrosis, focal necrosis and calcification of myocardial fibers in the epicardium and media layer, and purple-black calcium deposits, fibrous tissue and calcification in the necrotic area. lymphocytes. TSS treatment improved symptoms to a certain extent, suggesting that TSS has anti-inflammatory effects caused by CVB3 virus infection in vivo.

The in vitro antiviral effect study of TSS includes a time-of-addition test to determine its phase of action. The relative inhibition rate of the three administration methods on cell CPE: After virus entry treatment > drug pre-incubation treatment > simultaneous treatment of drug and virus. TSS can reduce CVB3 virus mRNA in a dose-dependent manner, which indicates that the mechanism of action of TSS is related to the inhibition of virus replication by entering cells. We detected that TSS can reduce the expression of inflammatory factors caused by CVB3 infection, suggesting that TSS also has anti-inflammatory effects in cells.

TLR3 has a critical role for the activation of immunity. TLR3 is able to recognize viral intermediate dsRNA through activation of TRIF, TRAF6, NF-κB. MAPK, AP-1, and IRF-3 drive the expression of proinflammatory cytokines^(13–15)^, that initiate the innate immune response to viruses leading to IFN expression^(16)^. We found that CVB3 infection caused elevated expression of TLR3 pathway related factors in mice and host HT-29 cell levels, and TSS could reduce the expression of TLR3 and its downstream cytokines, suggesting that TSS can play an anti-CVB3 virus role through TLR3 pathway.

Herein, a promising anti-CVB3 virus drug candidate TSS has shown viral infection therapeutic potential. The anti-CVB3 viral efficiency and molecular mechanism of TSS were preliminarily discussed. The host TLR3 inhibitory effect of TSS is the main mechanism of its anti-CVB3 viral activity. In addition, TSS can also improve immune organ index, enhance immune function, and downregulate the expression of some inflammatory factors. All these experimental results suggest multiple antiviral mechanisms of TSSs, which will be further investigated in our future work.

## Funding

This work was supported by the Sichuan provincial science and technology department (No. 2020YFS0370).

## AUTHOR CONTRIBUTIONS

Xiao-han Zheng designed the research and responsible for literature review and writing, Yuan Wang, Jing Zhou responsible for correction, Ming-mingYuan, Lei Zhang responsible for proofreading, literature review and correction. All authors read and approved thefinal manuscript.

## CONFLICT OF INTEREST

The authors declare that the research was conducted in the absence of any commercial or financial relationships that could be construed as a potential conflict of interest.

